# Density and temperature controlled fluid extraction in a bacterial biofilm is determined by poly-γ-glutamic acid production

**DOI:** 10.1101/2020.12.21.423644

**Authors:** Ryan J. Morris, Tetyana Sukhodub, Cait E. MacPhee, Nicola R. Stanley-Wall

## Abstract

A hallmark of microbial biofilms is the self-production of extracellular matrix that encases the cells resident within the community. The matrix provides protection from the environment, while spatial heterogeneity of expression influences the structural morphology and colony spreading dynamics. *Bacillus subtilis* is a model bacterial system used to uncover the regulatory pathways and key building blocks required for biofilm growth and development. Previous reports have suggested that poly-*γ*-glutamic acid (PGA) production is suppressed during biofilm formation and does not play a major role in biofilm morphology of the undomesticated isolate NCIB 3610. In this work we report on the observation of multiple travelling fronts that develop during the early stage of *B. subtilis* colony biofilm formation. We find the emergence of a highly motile population of bacteria that is facilitated by the extraction of fluid from the underlying agar substrate. Motility develops behind a moving front of fluid that propagates from the boundary of the biofilm towards the interior. The extent of proliferation is strongly modulated by the presence of extracellular polysaccharides (EPS). We trace the origin of this moving front of fluid to the production of PGA. We find that PGA production is correlated with higher temperatures, resulting in a mature biofilm morphology that is distinct from the biofilm architecture typically associated with *B. subtilis*. Our results suggest that *B. subtilis* NCIB 3610 produces distinct biofilm matrices in response to environmental conditions.

## Introduction

A common strategy employed by bacteria to mitigate stresses imposed by their environment is to co-exist in sessile communities known as biofilms. The transition from unicellular to multi-cellular life allows the residents to coordinate response to stimuli, share metabolic burdens ^1^, and protect against external attack by predators ^2,3^ or antimicrobial agents^4,5^. This behaviour is ubiquitous across the microbial world and a clear understanding of biofilm genesis, development, and maturation is important not only from a fundamental microbiological perspective but also due to their impact on many industrial, clinical, and biotechnological sectors. For example, biofilms act as sources for many chronic infections and their physical characteristics make them difficult to eradicate ^6,7^. This intransigence can also impact industrial processes, where biofilms may result in pipe blockages, induce corrosion, or contaminate products ^8–10^. While there are many negative consequences of biofilm formation, they play important roles in waste water treatment and other bioremediation processes ^11–14^.

*Bacillus subtilis* is a Gram-positive bacterium that has been extensively used as a model organism to illuminate the genetic regulation and molecular mechanisms of biofilm formation^15^. During biofilm formation, production of the extracellular matrix is an essential process that binds the cells together, provides protection from the external environment, and confers advantageous mechanical properties to the community. The primary components of the matrix produced by *B. subtilis* are the fibrillar protein TasA ^16^, the hydrophobin-like protein surfactant BslA^17–19^, and polysaccharides synthesized by products of the *epsA-O* operon ^20^. One of the principle regulatory pathways that controls the expression of these components is modulated by the transcription factor Spo0A. Moderate levels of phosphorylated Spo0A activates transcription of the *sinI-sinR* operon ^21,22^. SinR is a DNA-binding transcription factor that controls matrix production by interacting with the *epsA-O* and the *tapA-sipW-tasA* promoters ^23^. When SinR binds to its antagonist proteins (SinI and SlrR), repression is alleviated from these operons and biofilm formation can proceed^24,25^.

*B. subtilis* has several modes of active motility, two of which, swimming and swarming, are driven by the action of rotating flagella. Importantly, it has been shown that flagella-driven motility and biofilm phenotypes are bistable: cells can only express genes for motility or biofilm formation but not both at one time ^26,27^. Bi-stability allows for the emergence of phenotypic heterogeneity within a population of genetically identical cells. ^28^. This genetic bet-hedging provides a contingency for the community if and when environmental conditions change from their current state^28,29^.

In this work we report on the observation of multiple travelling fluid fronts that develop during the early stage of *B. subtilis* colony biofilm formation. We find the emergence of a highly motile population of bacteria that is facilitated by the extraction of fluid from the underlying agar substrate. Motility develops behind a moving front of fluid that propagates from the boundary of the colony towards the interior, and the extent of proliferation is modulated by a specific biofilm matrix component, the extracellular polysaccharide (EPS). We trace the origin of this moving front of fluid to the production of the polymer poly-*γ*-glutamic acid (PGA). We find the influx of fluid is dependent upon both bacterial density and environmental temperature. The temperature dependent production of PGA and the concomitant extraction of fluid can significantly impact the mature biofilm morphology, which diverges from the typical structure associated with *B. subtilis* NCIB 3610. Our results suggest that *B. subtilis* has the ability to produce an alternative extracellular matrix in response to adverse environmental conditions.

## Results

### Deposition and imaging of growing biofilms

At the beginning of each experiment, a 3 *μ*L suspension of *B. subtilis* cells (OD_600_ = 1) are deposited onto MSgg agar, a biofilm-promoting minimal media ^30^. After inoculation the droplet evaporates, which results in a ‘coffee ring’ deposition pattern: a higher density ring of bacteria accumulates at the edge of the initial droplet while the interior is more sparsely populated (Fig. 1A). This is caused by the differential rate of evaporation across the droplet; fluid evaporates more rapidly from the edge relative to the interior. This process drives capillary flows that transport bacteria from the interior to the boundary between the droplet and solid agar ^31^. Our initial experiments were performed using the wild-type isolate NCIB 3610 that was allowed to grow at 38°C while time-resolved images were collected. The images were captured by imaging through the agar substrate and we monitor the dynamics of growth on the underside of the biofilm (Fig. 1A). We acquired images for the initial ~ 6 hours of growth. Examples of images taken in this manner are seen in Fig. 1B-D. Here, the high density ‘coffee ring’ region is clearly identifiable by the large accumulation of bacteria near the outer boundary of the colony (Fig. 1B). This high density region of bacteria often appears to be multi-layered with a width typically ranging from 75-100 *μ*m.

**Figure 1.**
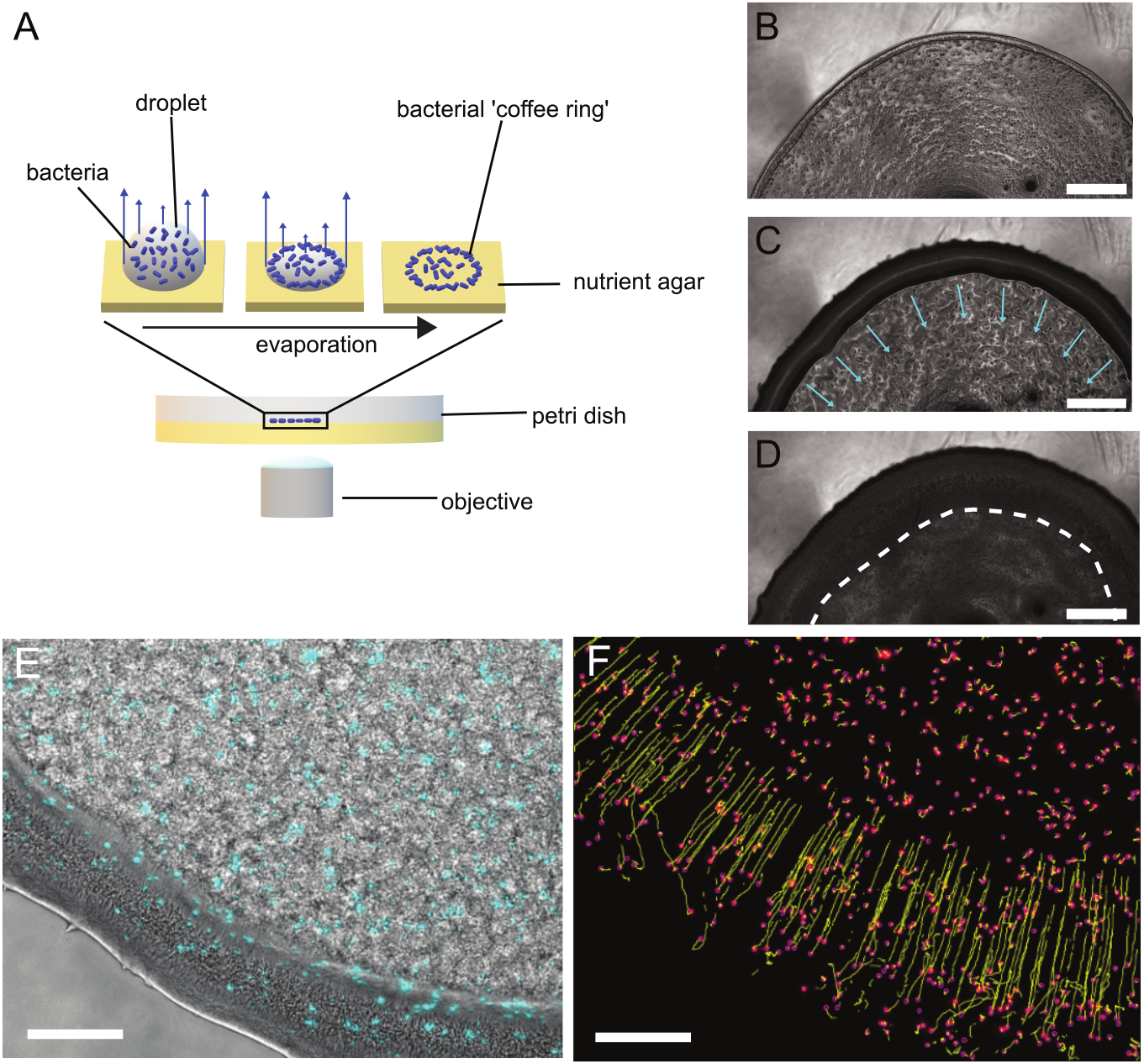
Schematic of the experiment and emergence of a travelling front of motile cells. (A) A 3 *μ*L droplet of a bacterial suspension is deposited onto the surface of MSgg agar and allowed to dry. Due to the differential rates of evaporation across the surface of the droplet (blue arrows), capillary flows are induced in the interior of the droplet. This allows for the transport of bacteria from the interior of the droplet to the edge where they become deposited. The effect is an accumulation of a higher density region of bacteria at the contact line between the droplet and solid agar. (B) This higher density region of bacteria can be seen at the edge of the colony. (C) After three hours of incubation and growth a darker region develops centered upon the edge of the emerging biofilm. We find that this zone is fluid and begins to move inwards towards the center of the colony (blue arrows). In this zone we observe active and motile cells. (D) After another two hours, the front stops moving inwards, indicated by the white dashed line, and the motility stops. Scale bars (B)-(D) are 500 *μ*m. Fluorescent beads were added to the suspension of bacteria and deposited with them at the beginning of the experiment; the beads are colored cyan. The ‘motility zone’ is identifiable as the darker region nearer the boundary of the colony (c.f. (C)). (F) Using particle tracking software, the beads are located (pink) and their motion tracked over time (yellow lines).The majority of beads move linearly at a constant rate towards the interior of the colony. Scale bars (E)-(F) 200 *μ*m.

### Emergence of a fluid flux that induces motility within the early biofilm

After this initial time we observe a zone, emerging from the high density region, that is visually different to the interior of the colony (c.f. the dark annulus at the edge compared to the lighter interior, Fig. 1C). This zone grows and concurrently moves inwards as a travelling front towards the interior of the colony (Movie S1). Upon closer inspection of the travelling front we observe a highly dense and motile population of bacteria. The motile cells in this confined region clearly exhibit coherent patterns of self-organised motion such as swirls and vortices, reminiscent of structures observed in active turbulent systems (Movie S2) ^32,33^. This behaviour implies that the cells are now in a fluid environment. The motility continues as this fluid zone grows, after which the motion rapidly stops (Fig. 1C) after travelling a distance of ≈ 400 *μ*m (Fig. 1D). This cessation of motility often also occurs as a propagating front, albeit travelling much faster than the initial propagation of motility (Movie S1, Movie S3) and predominantly moving from the interior of the colony back towards the coffee ring. Taken together, we infer that this moving front of motile cells represents a fluid flux into the biofilm.

Next, we added 2 *μ*m fluorescent beads to the suspension of bacteria that were deposited at the beginning of the experiment (Fig. 1E). Tracking the motion of the beads revealed that the beads coincident with the frontal zone are pushed along at a constant speed of ≈ 2.5*μ*m min^−1^ (Fig. 1F). Thus, the motion of this front has the ability to displace and mechanically push beads, but also cells, towards the center of the colony. A considerable portion of the beads remains embedded within the bacterial mass, but some become erratic in their movement. Our interpretation of this behaviour is that these beads are set in motion by the swimming action of the bacteria in a fluid environment. Interestingly, when we look at bead movement across the entire diameter of a colony, we also observe beads that move outwards radially from localized regions and this occurs simultaneously with the bead movement we see at the biofilm edge (Movie S4). Taking all data together, we infer that this moving front represents a fluid flux into the biofilm, and its emergence depends upon a high cell density.

### Extracellular polysaccharide production controls the spatial extent of the fluid flux

Since we observe the formation of a fluid annulus at the outer edge of the biofilm, this led us to investigate if the extracellular matrix controls and modulates the extent of the fluid flux. We hypothesise that the formation of an annulus is a result of extracellular matrix production within the central region of the biofilm that prevents further incursion of fluid. We performed the same experiment described in Fig. 1A using a strain that possesses a deletion of *sinR*, a key repressor of biofilm formation. Without *sinR*, the cells over-produce the extracellular matrix and the colony biofilm occupies a smaller footprint and is highly wrinkled (Fig. S1)^23^. In this case, we posit that the fluid annulus should be more confined if the matrix impacts the spatial extent of fluid flow into the biofilm. We measured the maximum distance that the fluid encroached into the interior of the biofilm relative to the outer edge of the colony (Fig. S2, Movie S5). We found that the distance the fluid travelled in the *sinR* strain was approximately 3 times less, on average, than the wild-type 3610 strain (Fig. 2). We conclude that extracellular matrix production can modulate the spatial extent of the fluid flux into the biofilm.

**Figure 2.**
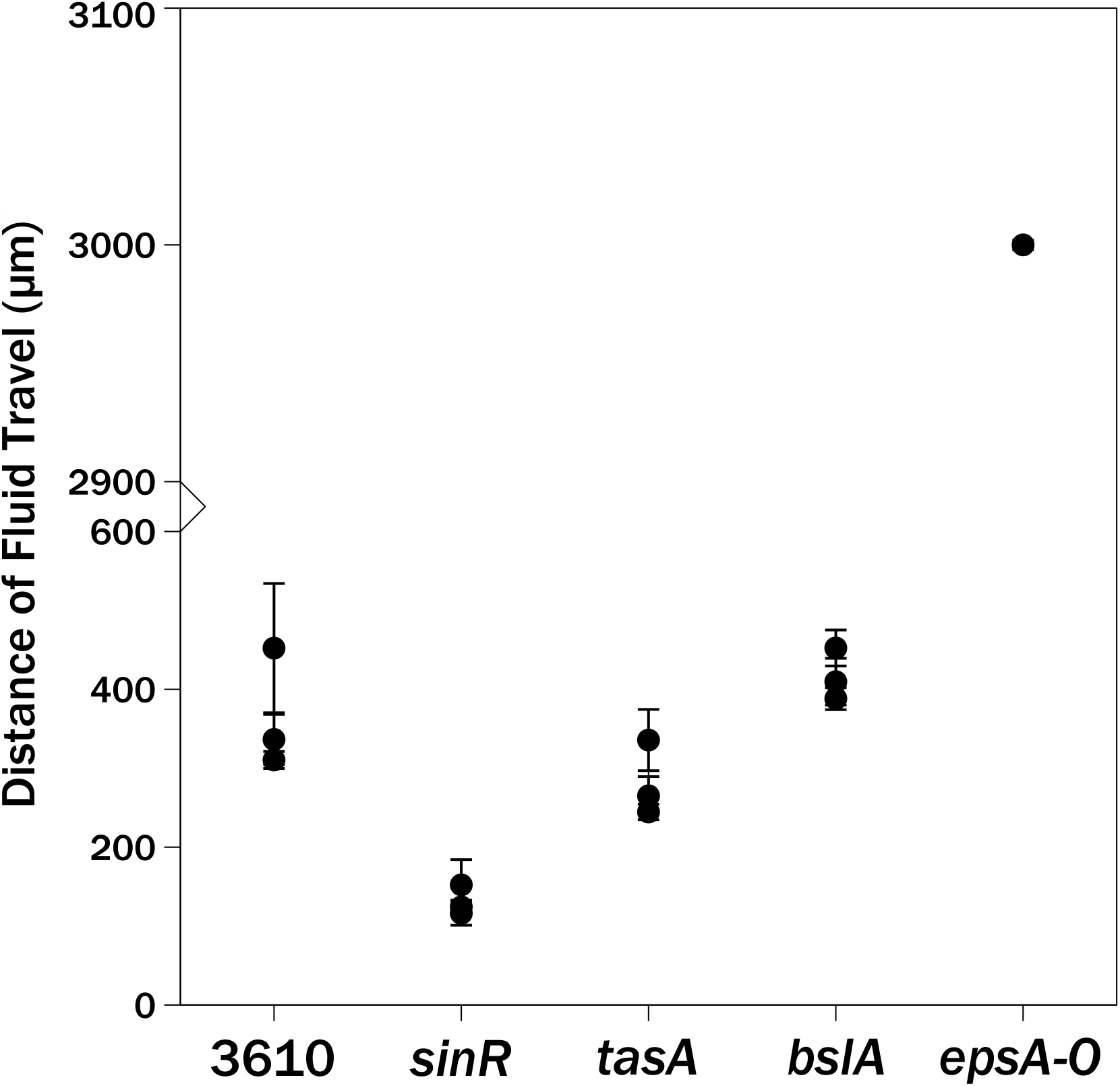
Matrix production can modify the spatial extent of fluid invasion. Plotted is the average distance of fluid travel as measured from the edge of the biofilm. The y-axis has been broken to show the differences between the 3610, *sinR*, *tasA*, and *bslA* strains while still representing the large distance of fluid travel found in the *epsA-O* strain. Each data point is the mean fluid travel distance averaged over 10 spatial points across an individual biofilm (N=3 for each strain). Error bars represent standard deviation.

Next, we wished to determine if one or more individual components of the matrix were important in controlling the fluid flux. We performed analogous experiments on strains that possessed gene deletions for BslA (*bslA*), TasA (*tasA*), and EPS (*epsA-O*), respectively and we again measured the distance of fluid travel. We found that the *tasA* and *bslA* strains were similar to the wild-type strain, albeit the fluid flux in the *tasA* strain was slightly less than the wild-type and *bslA* strains (Fig. 2, Fig. S2). In contrast, the strain that produced the greatest difference in behaviour compared to NCIB 3610 was the strain that does not produce EPS. In this strain, as the fluid propagates inwards it is not confined to an annulus but moves to the very center of the colony as evidenced by the very large fluid travel distance (Fig. 2, Movies S1, S6, S7). This fluid coverage resulted in motile cells being visible across the entirety of the colony. Again, the motility was highly dynamic and we observed vortex formation characteristic of active turbulence (Movie S6). The motility arrest front is even more apparent in this strain and, like the fluid propagation front, travels across the entirety of the colony (Movie S1, S7, S8). While the *tasA* and *bslA* strains showed very similar dynamics to the wild-type in the first 6-8 hours, the mature biofilms of these matrix mutants resulted in smaller and more unstructured colonies compared to the wild-type strain (Fig. S1). Additionally, the *epsA-O* mature biofilms were similarly small and morphologically unstructured (Fig. S1) implying under these conditions, the initial fluid flux does not drastically perturb the colony morphology, and matrix production is the important driver in determining mature biofilm structure. Taken together our data demonstrate that the EPS is the determinant that controls the degree by which the fluid flux invades the biofilm.

### Whole biofilm imaging of epsA-O strain

The pronounced fluid expansion and motility in the *epsA-O* strain allowed us to track the flux of fluid across the entirety of the biofilm in a time-resolved manner to learn more about the fluid extraction process. We noticed a finger-like instability develops as the fluid propagates inwards (Fig. 3A; Movies S7,S8). As the fluid pushes in towards the center of the colony the fluid front becomes unstable, forming increasingly large fingers over time (Fig. 3A,B). Such finger-like instabilities are well established in fluid dynamics. A well-known example is the Saffman-Taylor instability (also known as viscous fingering) that occurs when an unstable interface develops between a less viscous fluid displacing a fluid of higher viscosity^34^. Another way in which fingering morphologies can develop is through gradients in surface tension: Marangoni flows are induced when a fluid with lower surface tension flows towards regions of higher surface tension. Under appropriate conditions, a finger-like morphology can develop when an aqueous surfactant solution spreads on a thin-film of water ^35,36^.

**Figure 3.**
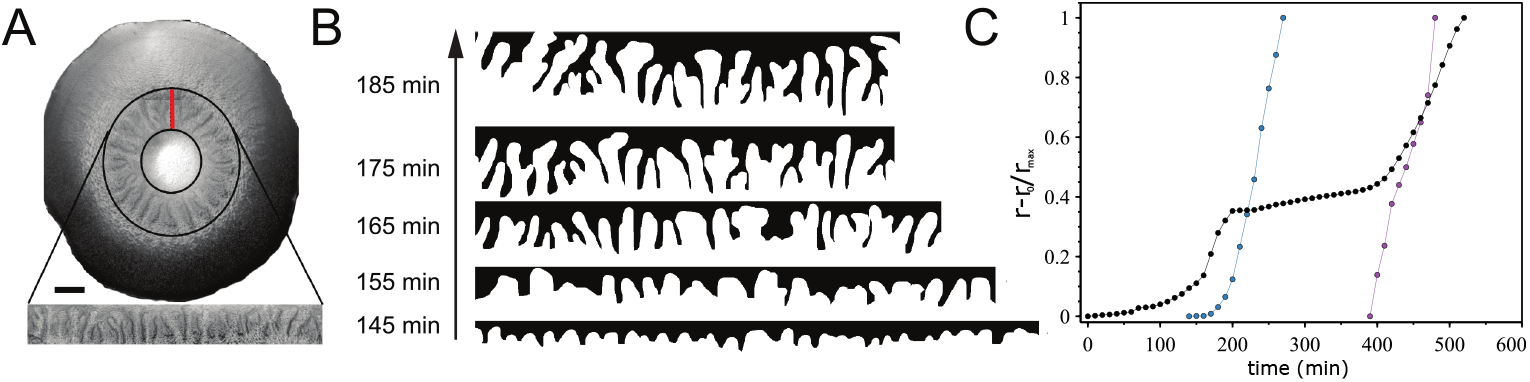
Large scale dynamics in polysaccharide-minus biofilms. (A) An example of a microscopy image of an entire *epsA-O* biofilm (scale bar is 500 *μ*m). Annular black region defines the area where we observe finger-like instabilities. This annular region is made linear for ease of visualization; a cut is made at the red line and the circular region is ‘unrolled’ to form a linear region. (B) Binarized images of the fingers over time for the biofilm in (A). (C) The normalized displacement as a function of time for the outer edge of the colony (black), fluid front (blue) and motility arrest front (purple). A plateau in the outer edge growth occurs coincidentally with the onset of fluid propagation. The growth recommences after motility becomes arrested. The blue curve corresponds to the time when we observe the fingers develop in (A).

Imaging the biofilms in this manner also allowed us to fully track three key features of the dynamics: (1) the distance travelled by the expanding outer edge of the colony, (2) the distance travelled by the fluid flux, and (3) the motility arrest front. All measurements were taken relative to the initial position of each feature. Additionally, the relative distances were normalized by their maximum value to allow comparison of their temporal evolution (Fig. 3B). Initially, there is a lag time before the outer edge of the colony begins to expand at a constant rate beyond the initial deposition position. We find that the outer edge growth slows down and plateaus at precisely the time that we see the fluid front begin to propagate. Moreover, the onset of the arrest front propagation directly coincides with the resumption of growth at the outer edge of the biofilm.

### Polyglutamic acid is the agent that induces the fluid flux into the biofilm

We wished to identify the molecule responsible for driving the fluid flux and subsequent motility and growth dynamics. Surfactin is a lipopeptide produced by *B. subtilis* that is a powerful biosurfactant ^37^ and a potent anti-microbial agent ^38^. Extracellular surfactin aids swarming and surface motility by lowering the surface tension of the fluid film and increasing the wettability of the substrate ^39,40^, as well as acting as a ‘one-way’ signalling molecule that induces the activation of extracellular matrix genes ^41^. Surfactin production in *B. subtilis* biofilms facilitates colony spreading ^40^ and is important in the osmotic extraction of fluid from the underlying agar substrate ^42^. Given these properties, we tested whether surfactin was the causative agent of the dynamics we observe. We used the *srfAA* and *srfAA epsA-O* deletion strains to compare the dynamics in both wild-type and EPS deficient backgrounds when surfactin could not be made. As a diagnostic for potential differences between the strains we again measure the relative displacement of the biofilm edge as a function of time. If the interior of the growing biofilm becomes fluidized we should see a plateau in the displacement curve as we saw in Fig. 3C. We found that when we measured the outer edge displacement for biofilms possessing these mutations, they both exhibited the characteristic ‘fluidization plateau’ (Fig. 4A). These data indicate that the *srfA* and *srfA epsA-O* strains did not abolish the fluid flux into the colony and led us to conclude that surfactin is not the causative agent of the fluid extraction.

**Figure 4.**
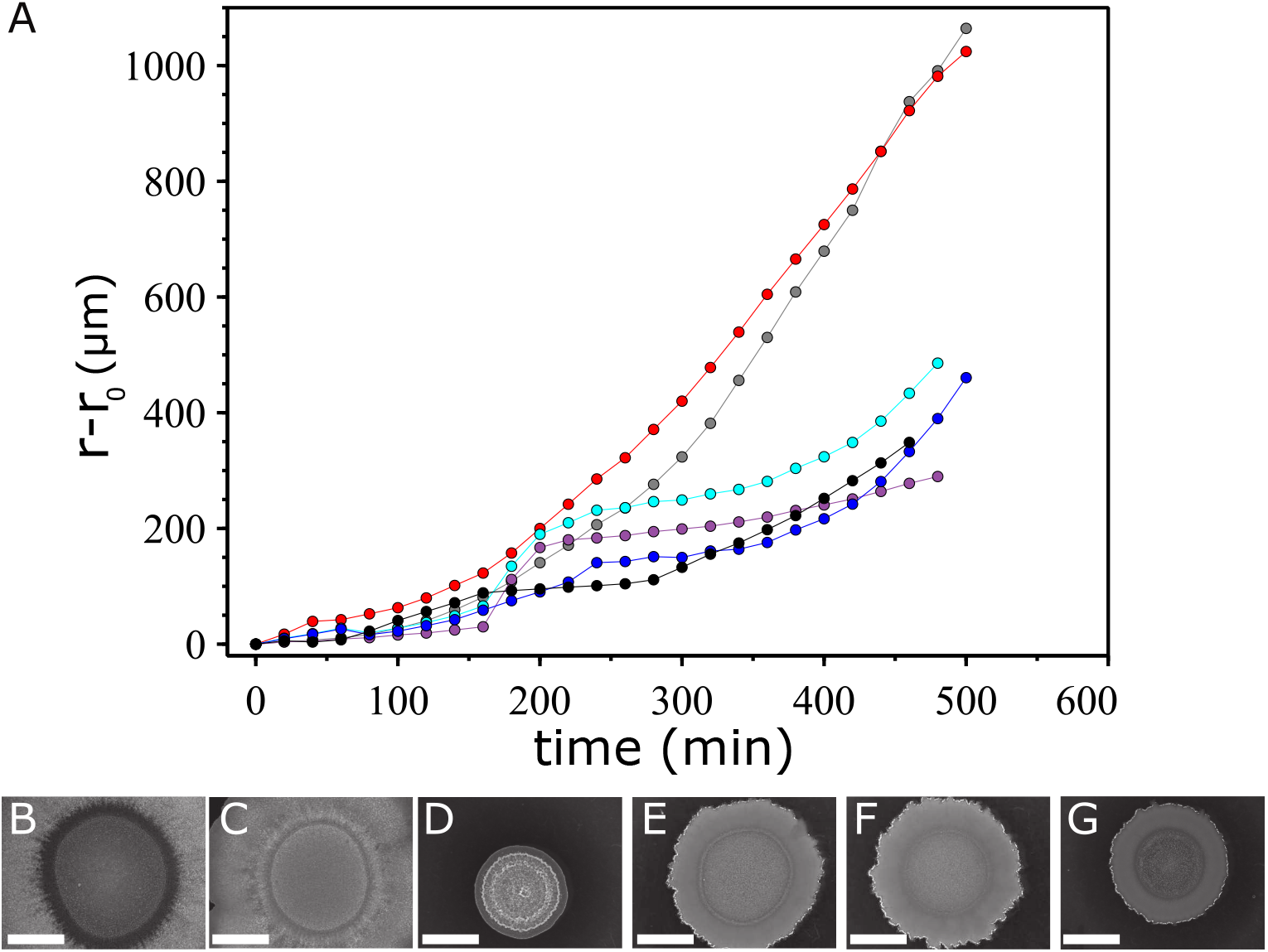
Fluid extraction is driven by PGA production. (A) Measurements of the outer edge displacement at 38°C show that NCIB 3610 (black), *epsA-O* (blue), the *srfAA* (purple), and *srfAA epsA-O* (cyan) strains possesses the characteristic ‘fluidization plateau’. The *pgsB* minus (red), and *pgsB epsA-O* minus strain (grey) that cannot produce PGA did not exhibit this behavior. Colony morphology after 48 hours incubation at 38°C of (B) NCIB 3610, (C) *pgsB* (D) *srfAA* (E) *epsA-O*, (F) *pgsB epsA-O*, and (G) *srfAA epsA-O*. Scale bar is 5 mm.

Poly-*γ*-glutamic acid is a naturally occurring biopolymer consisting of repeating units of L-glutamic acid, D-glutamic acid or both, and is primarily produced by species of *Bacillus* ^43,44^. PGA is another potential candidate due to its humectant properties ^45,46^ and its density dependent production ^47^. To test whether PGA was responsible for the fluid extraction and the characteristic growth curves, we produced two deletion strains that targeted *pgsB*, the gene encoding the essential synthetase required for PGA production^48^. One strain possessed a deletion of *pgsB*, while the other possessed a deletion of the gene *pgsB* in the *epsA-O* background. Again, we measured the edge displacement simultaneously for the wild-type, *epsA-O*, *pgsB*, and *pgsB epsA-O* strains. We observe a plateau in the edge displacement for the wild-type and *epsA-O* strains while we observe no change in the displacement rate after the lag time for the *pgsB* mutants (Fig. 4A). We also imaged the biofilm morphology after 48 hours of incubation and found that the wild-type and *pgsB* mutant were morphologically similar (Figs. 4B,C), and different to the strains unable to produce EPS which were likewise comparable (Figs. 4E,F). For completeness, we also imaged the surfactin mutant biofilms and found that the *srfA* strain was structured but occupied a small footprint while the double *srfA pgsB* mutant was less structured, similar to the other non-EPS producing strains (Figs. 4D,G). We therefore conclude that PGA is the molecular agent that extracts fluid from the substrate and drives the dynamics that we have thus far described. Moreover, the initial fluid flux into the biofilm does not appreciably alter the morphology, and matrix production still governs the structural phenotypes of the colonies.

### PGA expression is correlated with high temperature conditions

The discovery that PGA was involved in biofilm formation was unexpected, as while PGA has a role in biofilm formation in several isolates of *B. subtilis* ^49^ it had previously been excluded as a matrix molecule for NCIB 3610^50^. Its absence was reported to have no impact on colony biofilm structure^47,50^. We wanted to reconcile these divergent observations. We noted that our analysis was performed at 38°C while most other studies investigating *B. subtilis* biofilm formation are conducted at temperatures ranging from room temperature up to 30°C. We therefore hypothesised that PGA production is temperature-dependent. First, we repeated the experiments with wild-type, *epsA-O*, *pgsB*, and *pgsB epsA-O* strains at the additional temperatures of 30°C and 42°C (the highest temperature achievable in our microscopy incubator). At 30°C we did not observe any fluid flow into the colony (i.e. no front development and motility). Likewise, there was no characteristic plateau in the edge displacement curve for all strains, even the wild-type and *epsA-O* strains (Fig. 5A). This result implies that PGA is not produced at 30°C. At the higher temperatures of 38°C, and 42°C we do find a plateau in the edge displacement curves for the strains able to produce PGA, while no plateau is observed for the PGA deficient mutants (Fig. 4A and Fig. 5B).

**Figure 5.**
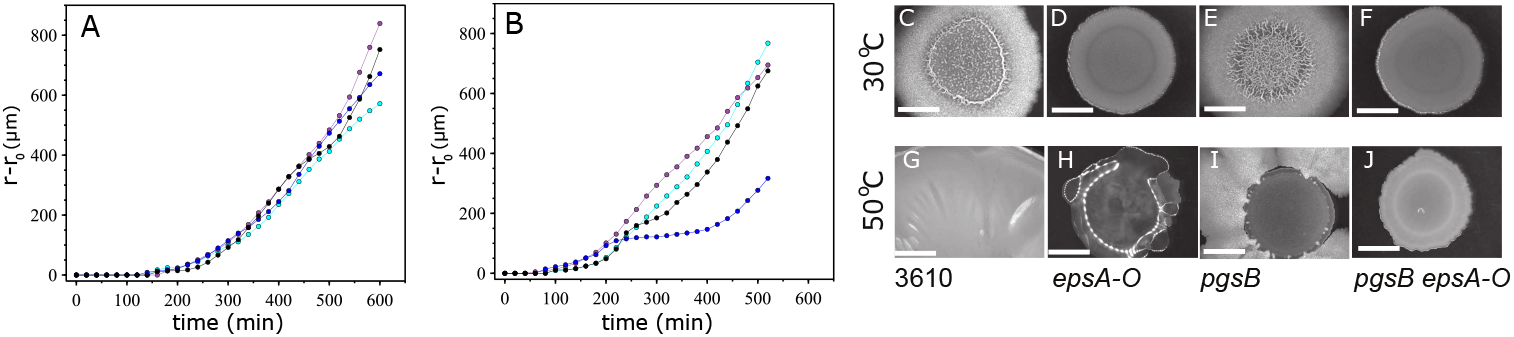
PGA production is temperature dependent. (A) Measurements of the outer edge displacement at 30°C and (B) 42°C for 3610 (black), *epsA-O* (blue), *pgsB* (magenta), *pgsB epsA-O* (cyan). Colony morphology after 48 hours incubation at (C)-(F) 30°C, and (G)-(J) 50°C. Scale bar is 5 mm.

Second, NCIB 3610 was found to form exceedingly mucoid colonies on MSgg agar plates at 50°C (Fig. 5G) that were amorphous and highly extended in shape and size. This further supports the idea that PGA production is temperature-dependent. The mucosity was at a level such that when the petri dish plate was inverted the biomass dropped onto the lid. We observed similar phenotypes for colonies unable to produce EPS and surfactin (Fig. 5H, Fig. S3). We also found the mucoid phenotype was directly linked with PGA production as, contrary to the NCIB 3610 parental strain, the *pgsB* deletion strains were entirely non-mucoid (Fig. 5I,J). We noted the *pgsB* strain appeared to develop cell flares extending from the edge of the initial colony biofilm footprint, which would be consistent with secondary mutations evolving.

When the same strains were examined at 30°C, no difference in the structure of the NCIB 3610 and *pgsB* colony biofilm architecture was observed (Fig. 5C,E), consistent with previous reports ^47,50^. Similarly, both the *epsA-O* and *pgsB epsA-O* mutants are morphologically comparable (Fig. 5D,F). This, in conjunction with results presented in Fig. 4B-G, implies that the determining factor for colony morphology at low to intermediate temperature conditions is the EPS component of the biofilm matrix. Conversely, at high temperatures, PGA dominates as the extracellular component that controls biofilm structure. We found that the matrix mutants containing deletions in *bslA*, *tasA*, and *epsA-O* formed similarly mucoid biofilms at 50°C (Fig. S3). Only the mutant possessing a deletion in *sinR*, while still mucoid, produced a biofilm with rugose structural features (Fig. S3). Collectively these data support the conclusion that PGA is produced in NCIB 3610 in both a density- and temperature-dependent manner and shows that biofilm architecture and structure can be dramatically influenced by the production of PGA.

## Discussion

### PGA production and motility

In this work we have shown that *B. subtilis* produces PGA that, due to its humectant properties, induces a fluid flux into a growing colony. This up-welling of fluid induces cell motility that, in these high density and confined conditions, results in turbulent dynamics. It has been well established that motile and biofilm matrix-producing cell states are mutually exclusive ^26,28^; any individual cell can be one but not both. It has also been demonstrated for both Gram-positive and Gram-negative species that active flagellated motility is often required for biofilm development and the role it plays is multi-fold ^51–53,53–56^. It has been shown that the presence of motile cells can dramatically influence the rate of colony expansion when motile and non-motile cells are co-cultured ^57^. Turbulent fluid flows generated by flagella-driven motility can drastically affect nutrient mixing ^58^, transport of passive ‘cargo’ ^59^, and may influence the formation of *B. subtilis* pellicle biofilms ^60^. It is not yet clear what function PGA-induced motility may play in *B. subtilis* biofilm development. However, it is intriguing that that the same transcriptional regulators required for motility^61^ are also necessary for PGA synthesis^47,62,63^. In our experiments we observe motility in the same spatial location where fluid influx and, by inference, PGA production occurs. Whether the motility we observe is simply a by-product of being in a fluid environment, or whether there is a direct evolutionary and functional link between PGA production and motility is still an outstanding question.

From our experiments it is clear that high temperatures induce the colony to withdraw a considerable volume of fluid from the agar substrate. Previous work has shown that PGA can confer protection to bacteria and increase survival under many different environmental stresses^64–67^. It is plausible that when a biofilm is subject to temperature stress, an over-production of PGA into the extracellular environment would both scavenge and retain moisture which otherwise may be lost through evaporation, thus preventing desiccation of the colony. Additionally, a potential advantage of PGA production in a changing environment would be the possibility of active or passive spreading, to escape and search for more suitable environs to colonize.

### Spatial heterogeneity in matrix production

Our experiments at intermediate temperatures (38°C) are suggestive of spatial structuring between PGA-producing cells at the edge of the colony biofilm, and matrix-producing cells in the middle (as has previously been reported ^68^), resulting in the annular confinement of fluid. Such spatial and temporal heterogeneity is a common feature in biofilms where the local microenvironment can strongly influence the phenotypic state of the cells. Phenotypic heterogeneity within a biofilm can be generated from variations in the chemical or physical environment ^69,70^, genotypic variations, and stochastic gene expression ^69^.

The heterogeneity in cell density imposed by the initial deposition conditions - and the formation of the ‘coffee ring’ - leads to the spatial pattern of fluid extraction that we observe. However, this is not the only means of generating density differences within a biofilm. Aggregates formed while growing in liquid culture can seed patches of higher cell density across the deposition footprint. Indeed, our results show that fluid invasion and, by inference, PGA production can occur in very local regions far away from the ‘coffee ring’ (Movie S4). This suggests that there is some critical density that determines whether a cell adopts a PGA producing state over a matrix-producing one. The spatial heterogeneity is transient at 38°C, and EPS becomes the dominant matrix component that determines the large-scale biofilm morphology. The fluid flux is ultimately stopped by the production of the EPS element of the matrix in the middle region of the biofilm (Fig. 6B). The motile cells that are close to the EPS-producing cells appear to stop moving, and a solid front advances from the middle of the colony outwards. One mechanism that could explain this abrupt transition involves the engagement of the “molecular clutch” EpsE, which binds to FliG and leads to motility arrest ^71^. However, we may rule this scenario out since we observe the propagation of the motility arrest front in the *epsA-O* strain. It remains an open question of what physical or biological mechanism governs this phenomenon.

**Figure 6.**
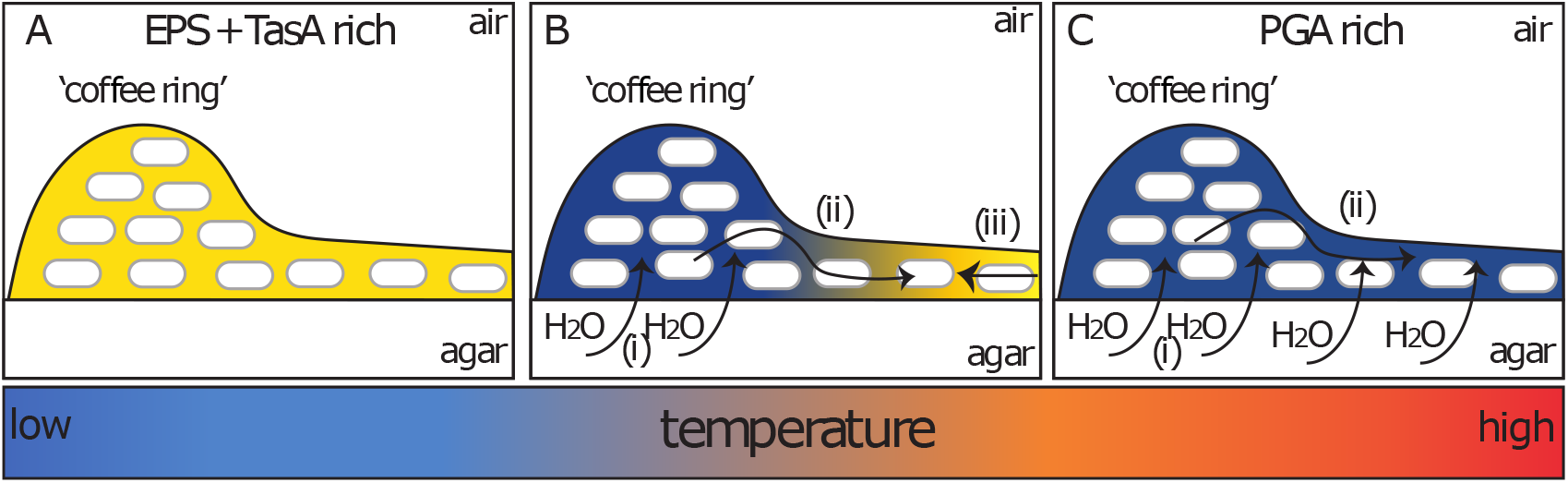
Schematic model of matrix production as a function of density and temperature. The ‘coffee ring’ is the initial region of higher density that is formed after deposition onto the agar surface. (A) Low temperatures produce a biofilm matrix rich in EPS and TasA (yellow). (B) Intermediate temperatures induces PGA production (blue) and (i) concomitant fluid extraction from the agar that originates in the high density region. (ii) The fluid propagates towards the center where (iii) EPS and TasA matrix production halts its advance. (C) High temperatures results in a PGA rich matrix that induces (i) fluid extraction that (ii) covers the entire biofilm resulting in a mucoid phenotype.

### Temperature dependent heterogeneity in matrix production

At low temperatures we did not observe the phenotypic heterogeneity that we observe at intermediate temperatures, presumably due to a lack of PGA production (Fig. 6A) despite the high cell density in the coffee ring. At high temperatures, biofilms are extraordinarily mucoid and lack any discernible structure typical of a *B. subtilis* biofilm (Fig. 6C). In either extreme, the local heterogeneities seem to matter little and density-dependent PGA production is superseded by a temperature-dependent pathway which impacts the entire biofilm.

When engineered to overproduce the TasA fibres and exopolysaccharide, by introduction of a mutation in *sinR*, the biofilms formed at high temperature (Fig. S3), were highly mucoid but also possessed structuring more typical of a wild-type *B. subtilis* biofilm. This implies that production of PGA can occur simultaneously with EPS/TasA, even if individual cells in the population commit to production of one or other product. This is also readily apparent at intermediate temperatures, where we infer that the two cell types have derived from the same population and are clearly present at the same time, albeit spatially separated.

Previous work has shown that strains possessing a deletion of *spo0A* result in biofilms that are mucoid and unstructured at low temperatures (30°C) and that this was due to PGA production^30,49^. Therefore, a *spo0A* mutant broadly mimics the wild-type biofilm phenotype that we observe under high temperature conditions. This implies that Spo0A may be the regulatory component that controls temperature dependent production of a PGA-rich or EPS/TasA-rich biofilm matrix.

Taken together, these results imply that *B. subtilis* has the ability to adapt different biofilm matrices with distinct properties to fit disparate environmental conditions. It is plausible that such a strategy is employed in natural environments as *B. subtilis* can grow at a wide range of temperatures: it is found in desert soils and within composts which can easily reach temperatures of 50°C. Until now, PGA was not thought to be an important factor as a matrix component in *B. subtilis* biofilms ^47^. However, it appears, under the right conditions, PGA can become an alternative matrix component with distinct structural and physical characteristics that may aid the biofilm to survive in high temperature conditions.

### Physical Consequences of PGA Production

The biofilm matrix of *B. subtilis* and *V. cholerae* has been modelled as a viscous hydrogel network that facilitates biofilm expansion via osmotic fluid influx ^72–75^. The localization of EPS production at the propagating boundary of a growing colony is thought to drive the outward expansion of the biofilm. Concomitant production of osmolytes stimulates fluid extraction that swells the matrix at the growing boundary, driving motion forward. In our experiments, PGA seems to have the opposite effect: colony expansion stalls due to the colony entering a ‘liquid-like’ state when fluid is extracted from the substrate. Previous work has shown that colony biofilm expansion is strongly governed by mechanical contact forces between neighboring cells and friction with the underlying substrate ^57,76,77^. In our experiments, expansion only recommences when the fluid environment dissipates, physical contacts are restored, and non-PGA matrix production begins. A question that remains is: how does colony expansion occur when PGA is the primary matrix component? Our experiments at high temperatures do show that the wild-type 3610 strain expands significantly beyond the initial deposition footprint. Strains that do not produce EPS or surfactin do not expand as much as the wild-type and this may hint that these components may still have a role to play in facilitating colony expansion (Fig. S3).

The traveling waves of fluid in the *epsA-O* deficient strain resulted in the appearance of finger-like structures as the wave propagated inwards. Such fingering instabilities can occur when a low viscosity fluid displaces one of a higher viscosity; the inverse situation will typically result in a stable interface. Curiously in our experiments, the fingers that we observe occur in the inverse configuration. Such inverse Saffman-Taylor instabilities can occur when there are wettable particles present that can adsorb to the air/fluid interface. This adsorption, due to interfacial energy minimization, can induce interfacial instabilities^78^. It is known that bacteria can accumulate at interfaces^79^, and *B. subtilis* can form floating biofilms at air/water interfaces. It is possible that bacteria coat and accumulate at the front of the incoming fluid wave, thus modifying the interfacial energetics and destabilising the interface between the fluid and the air at the interior of the colony. More work will need to be done to uncover the biological and physical mechanisms that cause this unusual phenomena and the possible benefit or function in ecological settings.

## Materials & Methods

### Growth conditions

*B. subtilis* strains were initially grown in LB medium (10 g NaCl, 5 g yeast extract, and 10 g tryptone per liter). Biofilms were grown on MSgg agar (5 mM potassium phosphate and 100 mM MOPs at pH 7.0 supplemented with 2 mM MgCl_2_, 700 *μ*M CaCl_2_, 50 *μ*M MnCl_2_, 50 *μ*M FeCl_3_, 1 *μ*M ZnCl_2_, 2 *μ*M thiamine, 0.5% v/v glycerol, 0.5% w/v glutamate, 1.5% w/v Select Agar (Invitrogen). When appropriate, antibiotics were used as required at the following concentrations: chloramphenicol at 5*μ*g ml^−1^, kanamycin at 25 *μ*g ml^−1^, spectinomycin at 100 *μ*g ml^−1^, and tetracycline at 10 *μ*g ml^−1^.

### Strain Construction

All strains used in this study are provided in Table S1. All *B. subtilis* strains used in this work are derived from the wild-type laboratory isolate NCIB 3610 and constructed using standard protocols. SPP1 phage transduction was used for transfer of genomic DNA from the donor strain into the recipient NCIB 3610, as described previously^80^.

### Biofilm Imaging and Analysis

*B. subtilis* strains were inoculated into 3 mL LB from a single colony grown on 1.5% w/v LB agar. The bacteria were allowed to grow at 30°C with 200 rpm orbital shaking until reaching an OD between 1.5-2. The cell culture was diluted to OD 1.0 in phosphate-buffered saline. A 3 *μ*L droplet of bacteria was deposited onto a 35 mm petri dish (Corning) containing MSgg agar. The droplets of bacteria were allowed to dry for 10 minutes. After drying, the petri dish was placed on a Nikon Ti inverted microscope that is temperature controlled. All microscopy images and movies were captured using a CoolSNAP HQ2 CCD camera controlled by *μ*Manager software. Brightfield movies and images of the biofilms were captured from the underside of the petri dish and are imaged through the agar. For the bead tracking experiments, 1 *μ*m diameter latex carboxylate-modified polystyrene yellow fluorescent beads (Sigma-Aldrich) were diluted into PBS from the stock solution in a 1:1000 ratio. 1 *μ*L of the working solution was added to the 1 mL of the diluted cell culture just prior to deposition. Movies were acquired in bright and epifluorescent channels (Nikon GFP fluorescent filter cube) and the beads were tracked using the ImageJ plugin TrackMate (v3.8.0)^81^. All image analysis was performed using the Fiji distribution of ImageJ. Whole biofilm microscopy images were captured as above but in multiple tiles that were stitched together using the ‘Pairwise Stitching’ plug-in. The images of the fingering instabilities were generated by first using the ‘straighten’ tool to transform a circular to a linear region. The default ImageJ threshold method was applied to binarize the images. In cases where the intensity varied across the image, the image was partitioned into regions of similar intensity and then thresholding was performed. The ‘edge finding’ tool was used to locate the edges of the thresholded images. After edges were identified the interior was filled to form a representation of the fingers. If the image was partitioned, the image was stitched together using the stitching tool in ImageJ. Measurement of fluid travel distance was manually tracked in ImageJ and the mean displacement was averaged over 10 separate measurements for each experiment. For each strain studied, three separate experiments were performed. Edge and front displacement measurements were similarly performed in ImageJ by manually tracking the movement of the front over successive frames. Displacements were always measured relative to the initial position of each feature. Images of whole biofilms were obtained using a Leica MZ16 stereoscope as described previously ^17^.

## Acknowledgements

We thank M. Porter and F. Davidson for their helpful comments. Work in the NSW and CEM laboratories is funded by the Biotechnology and Biological Science Research Council (BBSRC) [BB/P001335/1, BB/R012415/1].

