## Supplementary Information for "Density and temperature controlled fluid extraction in a bacterial biofilm is determined by poly-γ-glutamic acid production"

### FIGURES

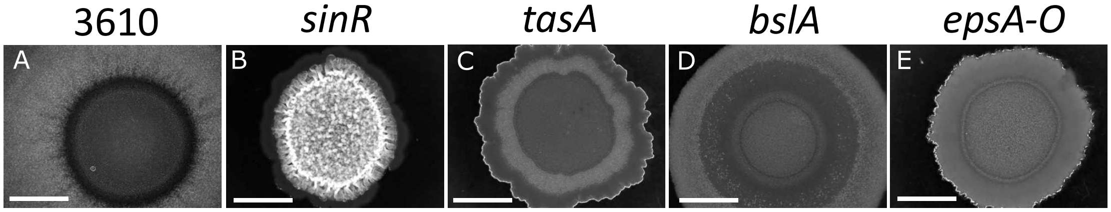


**Figure 1.** Representative colony biofilm morphology of matrix deficient strains at 38°C after 48 hours incubation. Scale bars are 5 mm.

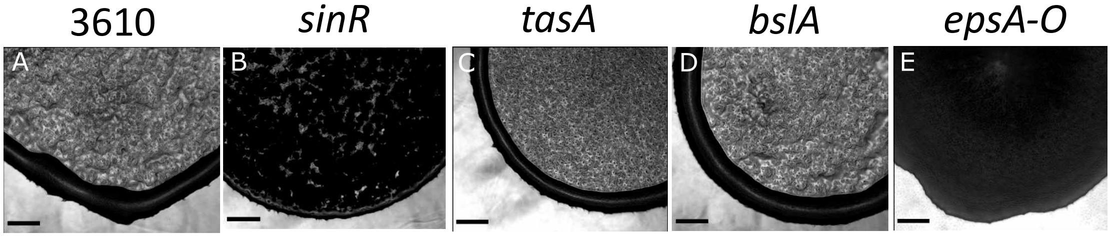


**Figure 2.** Microscopy images of matrix deficient strains taken from below through the agar at 38°C. Scale bar is 200 $\mu$m

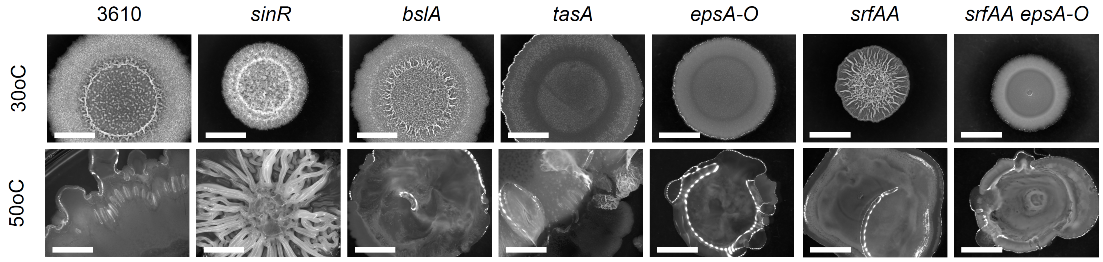


**Figure 3.** Representative colony biofilm morphology of matrix and / or surfactin deficient strains at 30°C and 50°C after 48 hours incubation. Scale bars are 5 mm.

### TABLES

| Strain | Genotype | Reference / Construction |
|---|---|---|
| NCIB 3610 | wild-type prototroph | BGSC |
| NRS2415 | Δ*tasA*::spc | [1] |
| NRS2450 | Δ*epsA-O*::tet | [2] |
| NRS2097 | Δ*bslA*::cml | [1] |
| NRS1859 | Δ*sinR*::kan | [3] |
| NRS6962 | Δ*srfAA*::kan Δ*epsA-O*::tet | NRS2450 → NRS6962 |
| NRS6958 | Δ*srfAA*::kan | BKK03480 → NCIB 3610 |
| NRS7014 | Δ*pgsB*::spc | BAL1811 → NCIB 3610 |
| NRS7015 | Δ*pgsB*::spc Δ*epsA-O*::tet | BAL1811 → NRS2450 |
| BAL1811 | JH542 trpC2 pheA1 Δ*pgsB*::spc | [4] |
| BKK03480 | Δ*srfAA*::kan | [5] |

**Table 1.** Table of strains used in this work. Drug resistance cassettes are indicated as follows: cml, chloramphenicol resistance; kan, kanamycin resistance; tet, tetracycline resistance; and spc, spectinomycin resistance. BGSC represents the Bacillus genetic stock center. The direction of strain construction is indicated with phage SPP1 (→) recipient strain.

### MOVIE CAPTIONS

**Movie S1** Moving fronts in growing *B.subtilis* biofilms. The lower colony is 3610 wild-type; upper colony is the *epsA-O* strain. Frame rate is 1 frame $min^{-1}$. One can see the 'coffee ring' of higher density at the outer edge of the colonies at the start of the experiment. In both cases, the fluid originates from this region. The speckling that is visible after fluid extraction corresponds to motile cells. The fluid front moves in towards the center of each colony. After some time the motility comes to an end. The 'motility arrest front' is very apparent in the *epsA-O* strain.

**Movie S2** Turbulent motility in NCIB 3610.

**Movie S3** Motility arrest front in NCIB 3610 strain.

**Movie S4** Radial bead movement occurs throughout the colony and occurs simultaneously with the bead movement at the outer edge of the biofilm. Circles highlight regions of radial bead movement.

**Movie S5** Closeup of the fluid invasion for the *sinR* strain (see Fig S2). Due to the large amount of matrix being produced, fluid entering the biofilm is obscured. However, one can see the 'smoothing' of the edge of the biofilm which is indicative of the fluid entering the biofilm (c.f. Movie S1). Scale bar is 100 $\mu$m. Frame rate is 1 frame every 2 minutes.

**Movie S6** Turbulent motility dynamics in the *epsA-O* strain. The edge of the biofilm is located on the right edge of the frame.

**Movie S7** Example of *epsA-O* dynamics where we observe fingering and a clear 'motility arrest front' that propagates across the colony. The fluid front begins to propagate at 176 minutes. The fingering instability becomes visible at 226 minutes and persists until they reach the center at around 290 minutes. The motility arrest front begins near the outer edge around 380 minutes and propagates inwards towards the center. Note the outer edge growth becomes notable after approximately 400 minutes.

**Movie S8** Example of fingering instability observed in the *epsA-O* strain. Note the movie was made by stitching two movies together, hence the linear visual artifact in the top third of the biofilm image.
